# Improving the Generation and Selection of Virtual Populations in Quantitative Systems Pharmacology Models

**DOI:** 10.1101/196089

**Authors:** Theodore R. Rieger, Richard J. Allen, Lukas Bystricky, Yuzhou Chen, Glen Wright Colopy, Yifan Cui, Angelica Gonzalez, Yifei Liu, R. D. White, R. A. Everett, H. T. Banks, Cynthia J. Musante

## Abstract

Quantitative systems pharmacology (QSP) models aim to describe mechanistically the pathophysiology of disease and predict the effects of therapies on that disease. For most drug development applications, it is important to predict not only the mean response to an intervention but also the distribution of responses, due to inter-patient variability. Given the necessary complexity of QSP models, and the sparsity of relevant human data, the parameters of QSP models are often not well determined. One approach to overcome these limitations is to develop alternative virtual patients (VPs) and virtual populations (Vpops), which allow for the exploration of parametric uncertainty and reproduce inter-patient variability in response to perturbation. Here we evaluated approaches to improve the efficiency of generating Vpops. We aimed to generate Vpops without sacrificing diversity of the VPs’ pathophysiologies and phenotypes. To do this, we built upon a previously published approach (Allen, Rieger et al. 2016) by (a) incorporating alternative optimization algorithms (genetic algorithm and Metropolis-Hastings) or alternatively (b) augmenting the optimized objective function. Each method improved the baseline algorithm by requiring significantly fewer plausible patients (precursors to VPs) to create a reasonable Vpop. #ddct #qsp

## 3 Introduction

Physiologically based mathematical models are often used to describe and predict the response of a patient to an existing therapy or novel agent. These models, frequently referred to as quantitative systems pharmacology (QSP) models, are used to simulate clinical trials in drug development (Musante, Ramanujan et al. 2017). In these applications, it is important that they not only capture the mean patient response to treatment but also inter-patient variability and how that variability may evolve over time. In addition, due to the novel nature of many therapies; the complexity of human physiology; and generally limited human data, QSP models are rarely fully determined by data. One approach to these challenges is to develop alternate parameter sets to capture the variability in the real clinical trial population and sample as much uncertainty in the model parameters as possible (Gadkar, Budha et al. 2014, Hallow, Lo et al. 2014, van de Pas, Rullmann et al. 2014).

Previously, we published an algorithm for the generation and selection of these alternative value sets (Allen, Rieger et al. 2016). The algorithm used simulated annealing (SA) to generate as large a population of “plausible patients” (PPs) as was practical. SA was used with a cost functional that optimized solutions to be biologically feasible. These PPs, which we call a “plausible population”, were termed plausible since each generated parameter set simulated a patient that was physiologically reasonable, and could be in a clinical trial, but there was not yet any selection for how *likely* it was for that patient to have been in a particular clinical trial. We then used our novel selection technique to choose those patients from the plausible population that most resembled a desired clinical population. These selected patients were then termed virtual patients (VPs), and as a collection they were called a virtual population (Vpop).

Since the original algorithm created a plausible population that was naÏve to the targeted Vpop distribution, significant computational effort was expended in generating PPs in unlikely regions of the target distribution. Here we propose alternative algorithms to improve the generation of the plausible population for more efficient generation of the Vpop. The common approach for each newly tested algorithms is to use information about the target distribution in generating the plausible population. We explored this idea in three ways:

1. Nested simulated annealing (NSA), which augments the SA method by targeting PP generation using the probability density function of the target distribution;
2. A genetic algorithm (GA), which iteratively builds a plausible population according to a fitness function defined by the desired distribution; and lastly
3. A Metropolis-Hastings (MH) inspired importance-sampling technique.

Results of the original SA method were re-generated, for direct comparison to each of the three new approaches.

## 4 Methods

This current work is the evaluation of three approaches for the generation of Vpops that match distributions of clinical cohorts or populations. The general flow for our algorithm is (Figure 1):

**Figure 1.**
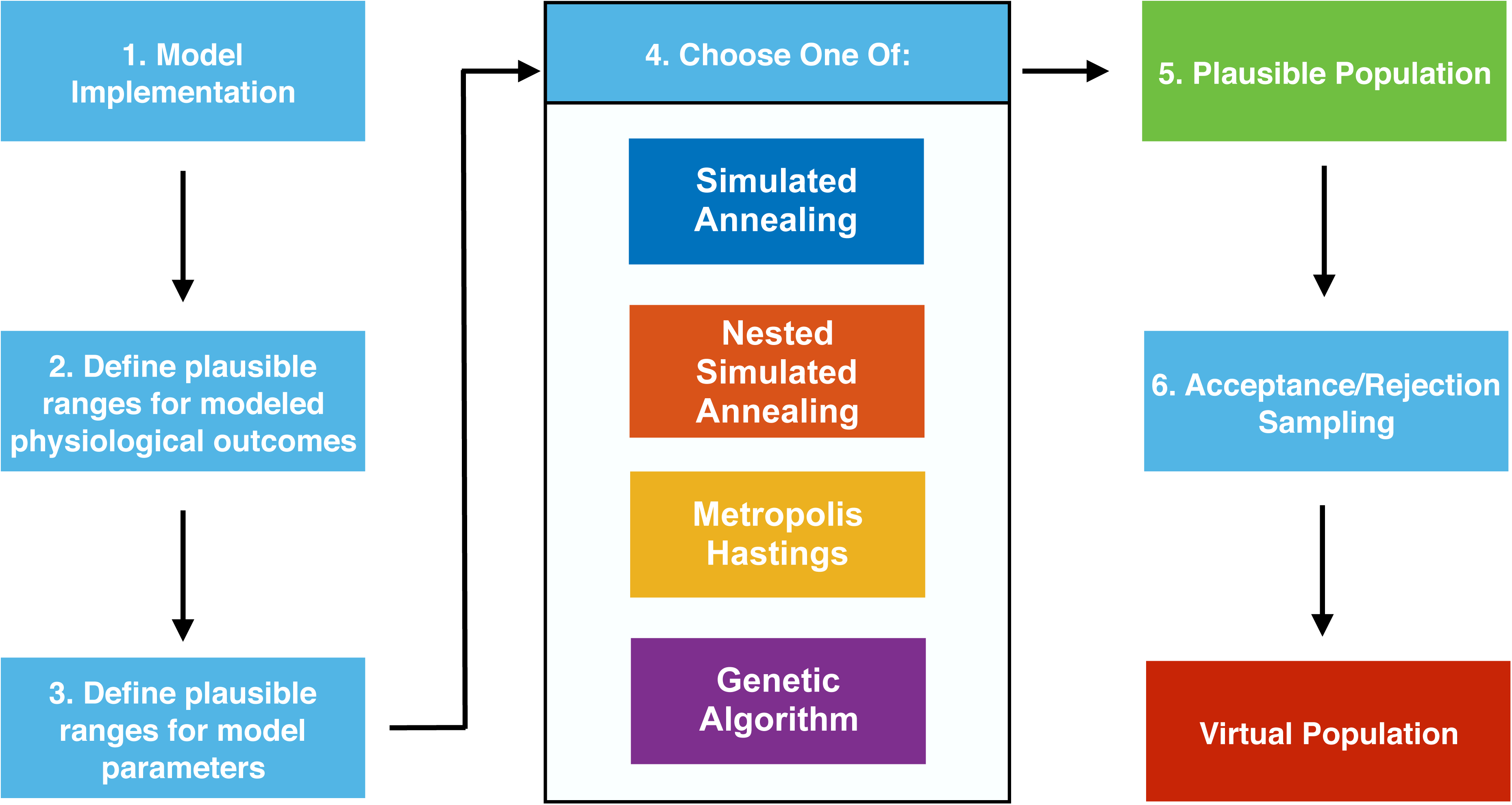
Flowchart of algorithm. Flowchart of algorithm. The initial problem setup is shared by each of the methods attempted, where biologically plausible ranges are placed on model parameters and states. Once the problem is setup, the generation of plausible patients is carried out by one of four algorithms. Post-plausible-patient generation, each algorithm follows the same acceptance/rejection sampling steps to create the virtual population.

1. Implement an ordinary differential equation (ODE) model that describes the biological system of interest;
2. For each state (variable) in the model, define a lower and upper limit for assessing if a steady state solution is plausible (all states between lower and upper limits) or not;
3. For each parameter of the model (e.g., rate constants, Michaelis-Menten constants), also define a plausible lower and upper limit for the search algorithms;
4. Optimize, using one of four algorithms, for solutions of the model that are PPs;
5. Collect the PPs generated by the optimization into a plausible population, terminating the search for PPs when the optimization achieves a preset number of PPs in the plausible population;
6. Perform acceptance/rejection sampling on the plausible population to select the VPs from the PPs that allow us to match the statistics of the target clinical population.

This section is organized to describe these three methods (NSA, modified GA, and modified MH), and how to apply them to generate VPs. This is followed by a description of how the results were analyzed, including a novel metric to quantify the uniqueness of a collection of parameter sets.

### 4.1 Mathematical Model and Data

Following our previous approach, we tested our proposed methods using a published ODE model of lipoprotein metabolism (van de Pas, Woutersen et al. 2012). In brief, the van de Pas model is a model of cholesterol production by the liver and its transit through the plasma. The ODE cholesterol model has nine equations, or state variables (Supplementary Table S1). These state variables correspond to the mass or concentrations of species within particular compartments (e.g., liver, plasma, peripheral tissues). The focus or primary outputs of the model are calculating levels of high-density lipoprotein cholesterol (HDL_c_) and “non-high-density lipoprotein cholesterol”, which we assumed to be equivalent to low-density lipoprotein cholesterol (LDL_c_). Thus the parameters of the ODE model we changed in the global optimization methods were the rate constants of the mass action model (e.g., production, reaction, clearance constants, see Supplementary Table S2). For the present work of creating baseline PPs and VPs, we only used the ODE model to simulate physiologically reasonable patients at steady state and were not concerned with the transient changes of the model.

While the ODE model has nine states, not all of them are frequently collected in clinical trials. The outputs of the model we matched to the statistics of human clinical data through our Vpop selection algorithm were: HDL_c_, LDL_c_, and total cholesterol (TC). As in original paper, for our reference population to match we used the National Health and Nutrition Examination Survey dataset (NHANES 2011-2012). The NHANES dataset contained fasting values for plasma cholesterol levels in 2,942 patients. The data was well represented by a joint lognormal distribution (Supplementary Figure S1).

### 4.2 Summary of Original SA Algorithm (Baseline Comparator)

In (Allen, Rieger et al. 2016) we optimized the steady-state solutions, *x*^*^, to fall within biologically reasonable ranges rather than to a specific point using the cost functional

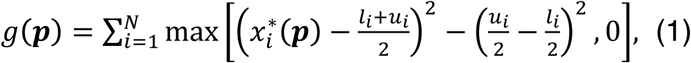

where *N*is the number of states of the model (N = 9 for van de Pas et al.), and *l_i_* and *u_i_* represent the biologically plausible lower and upper bounds for the *i^th^* state variable’s steady-state solution (see Supplementary Table S1). Note that if 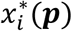 is within the plausible biological bounds, then *g****(p)*** = 0, and the parameter set ***p*** is called a PP. SA iteratively proposes values of ***p*** until *g****(p)*** = 0. A new initial ***p*** is then generated, and the SA optimization is repeated until a plausible population is created.

### 4.3 NSA Algorithm

The NSA algorithm incorporates information about the target data distribution in the process of generating the plausible population. By modifying the cost functional, equation (1), we can include information about the desired distribution, thus decreasing the number of PPs required per VP. The joint distribution of NHANES patients LDL_c_, HDL_c_, and TC is well approximated by a multivariate lognormal distribution, with a probability density function that is roughly ellipsoidal in iso-density lines. The hypercube in which *g****(p)*** = 0 in equation 1, was changed to an ellipsoidal shape, in the dimensions 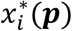 that correspond to LDL_c_, HDL_c_, and TC. The edge of this ellipsoid corresponds to the least-probable value (with respect to the multivariate log-normal) for which *g****(p)*** = 0.

However, the modified g(p) makes no distinction between low-probability solutions, at the edge of the ellipse, and high-probability solution, near the centroid of the ellipsoid. Hence, to fully use information from the target distribution, we considered multiple nested ellipsoids, relegating acceptable PPs according to whether they achieve sufficiently high probability density with respect to this target distribution. From this, the modified g(p) controls the number of PPs generated in each ellipse.

Note that while biological upper and lower bounds exist for all of the nine model outputs, we only have distributional information regarding LDL_c_, HDL_c_, and TC. As a result, the modified cost function still achieves a minimum when the remaining six dimensions falls within the hypercube of biologically plausible solutions.

We use the following equation for the surface of an ellipsoid that will encompass the data:

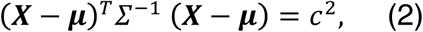

where ***X*** is a 3 × 1 vector representing a point in the log-scaled vector of [LDL_c_, HDL_c_, and TC], and ***μ*** and *Σ* are maximum likelihood estimates of the mean and covariance matrix of the multivariate normal distribution, and *c*^2^ controls the extreme point of the ellipsoid. By letting ***X*** be the data point furthest from the mean in (2), we can explicitly calculate the minimum value for *c*^2^ that allows the data to be encompassed by the ellipsoid.

First consider only one ellipsoid region. Then the modified cost functional is

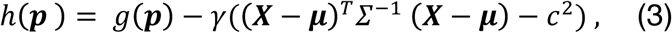

where *g****(p)*** is given by equation (1) for N = 6, and is calculated for the 6 state variables for which we do not have distributional information. We can interpret this cost functional as forcing these 6 state variables to be within plausible biological bounds while forcing the remaining 3 log-scaled observables to be within the smallest ellipsoid region that encompasses the clinical data. Notice we have an additional parameter, *γ*, which is a scaling factor that is a free parameter of the method for balancing the weight of the ellipsoid term on the cost functional. A value of *γ* = 0.1 was used for all simulations (Rieger and Allen 2017).

Now consider several nested ellipsoid regions. The cost functional for each nested ellipsoid is given in (3) but with modified *c*^2^ values in order to control the distribution of the plausible population. The number of nested ellipsoids, *R*, is another free parameter of this method. We denote the ellipsoids as *E*_1_ ⊂ *E*_2_ ⊂…⊂ *E_R_*, with the *k^th^* ellipsoid centered at the mean ***μ*** and defined as *E_k_* = **{*X*:(*X*** − 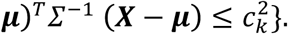

We choose the *c_k_* values such that a *k/R* proportion of the observations are within the *k^th^* ellipsoid:

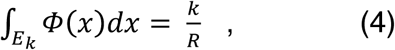

where *𝛷(x)* is the multivariate normal distribution, and *k* = 1… *R*. We find an approximate solution, *c_k_*, to this integral, by using a Monte Carlo approach. Equivalently,

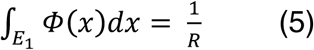

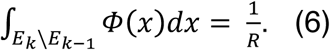

For this method, we sequentially populate each ellipsoid uniformly such that the final plausible population approximates the distribution for the target patient distribution. We therefore need to calculate how many PPs are required for each ellipsoid, given a desired total number of PPs. Define *q_k_* as the proportion of the total plausible population within the *k^th^* ellipsoid. Then, since we assume the data are approximately uniformly distributed throughout each ellipsoid, we want to solve a system of R equations for each *q_k_* obtained by solving

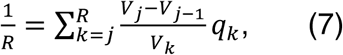

for *j* = 1,…, *R*. Where *V_j_* is the volume of the *j^th^* ellipsoid. We define *V*_0_ = 0. Note that 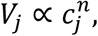 where *n* is the dimension of the multi-dimensional distribution. Then, equation (7) can be re-written as

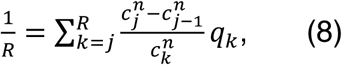

which can be solved recursively for the *q_k_* (starting with *k* = *R*, and defining 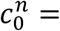 0).

Alternatively, since the NHANES data provided individual level data, the target distributions *c_k_* (and *q_k_*) were calculated empirically, see source code (Rieger and Allen 2017).

With the ellipsoids defined we can generate the plausible population by randomly generating a parameter set in the ranges we defined from a uniform distribution. Then, for a pre-defined plausible population of size *m*, for *l* = 1 to *m*, we identify a PP by minimizing:

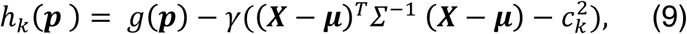

where *g****(p)*** is given by equation (1) for N = 6, and is calculated for the 6 state variables for which we do not have distributional information. See Supplementary Text 1 for further details.

This method requires choosing the number of nested ellipsoids. The greater the number of ellipsoids, the closer the distribution of the plausible population will match that of the clinical population, thus reducing the number of PPs required per VP. However, an increase in the number of ellipsoids increases computation time for the plausible population. In this case, we used five ellipsoids. We then use our rejection sampling method to select the VPs from the plausible population.

### 4.4 Modified GA

A commonly used population-based approach for optimizing nonlinear models is a genetic algorithm (GA), (Golberg 1989, Conn, Gould et al. 1991). For our problem, we created our plausible population using MATLAB’s *ga* function (MATLAB 2016). This algorithm first creates an initial population, where each patient is drawn from a uniform distribution. The algorithm then assigns a fitness value to each patient using a cost (fitness) functional. Similar to the NSA method, we modify equation (1) by incorporating information about the desired distribution. Specifically, the cost functional is given by

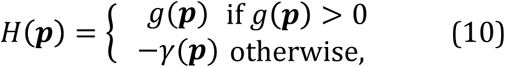

where *γ****(p)*** is the likelihood of a given log-scaled observable value ***X*** given by

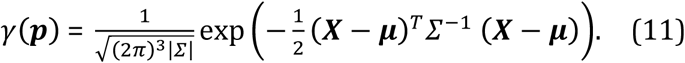

This cost functional can be interpreted as forcing all state variables to be within plausible biological bounds and additionally assessing a penalty as the log-scaled observables deviate from ***μ***.

As the algorithm progresses, children are created for each generation; those with a cost functional value *H****(p)*** < 0 become PPs. Since many PPs are created each generation, there is no way to specify the exact number of PPs generated. Thus we must preset the minimum number of PPs desired, but in practice we tended to generate slightly more than sought (see Supplementary Code). Once the plausible population is created, we use rejection sampling to determine the Vpop from the plausible population.

### 4.5 Modified MH Algorithm

The Metropolis-Hastings (MH) algorithm can be used to approximate a desired distribution, for a review see (Robert 2015). To apply the MH approach here we need to modify the algorithm. To see why, consider the original algorithm. Let *𝜋**(p)*** be our desired target multivariate probability distribution for the vector ***p***. The MH algorithm generates a sequence of ***p***, such that the distribution of this sequence,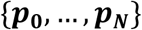 converges to *𝜋**(p)*** as *samplings →* ∞. Let *Q**(p, q)*** be some symmetric proposal distribution, which is interpreted as generating a proposed value ***q*** from *Q****(p, q)*** when the process is at value **p**. Then the original MH algorithm is as follows:

1. Generate an initial vector ***p*_0_**, set *i* = 1.
2. Generate a proposed vector ***p^*^~** Q**(P_i−1_, p^*^)***.
3. Calculate the probability ***p***^*^ is accepted, *α* = min 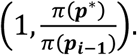
4. Generate *Y~U*(0,1), if *y ≤ α*, set ***p_i_*** = ***p***^*^. If *Y > α*, set ***p_i_*** = ***p_i−1_***.
5. Repeat steps 2 to 4 for *i* = 1,…, *number of samples* to collect {***p_0_,…, p_N_***} as a sampling from the target distribution.

This MH algorithm approximates the target distribution *𝜋**(p)*** by randomly sampling from it. At first glance, this approach appears immediately applicable to the problem at hand and will generate a plausible population that will converge to the Vpop as *N* → ∞. However, this algorithm requires modification because we do not know *𝜋**(p)** a priori*; i.e., we do not know how the parameters sets should be distributed such that the model, when simulated using those parameters, matches the data.

We rewrite our target distribution as *T**(X_p_)***, where ***X*** is the observable outcomes generated by the model *M*, using a parameter set (which in this case is in the log-space, so ***X_p_*** = log *M****(p)***). Then our algorithm becomes

1. Generate an initial vector ***p*****_0_**, set *i* = 1.
2. Generate a proposed vector ***p***^*^~ *Q****(p_i−1_, p^*^)***, write *Q_m_****(p_i−1_, p^*^)*** = *M(Q**(p_i−1_, p^*^)**)*.
3. Calculate the probability ***p***^*^ is accepted, *α* = min 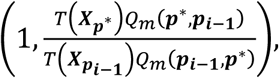 assume *α* ≅ min 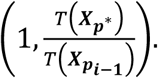
4. Generate *y~U*(0,1), if *y* ≤ *α*, set ***p_i_*** = ***p***^*^. If *y > α*, set ***p_i_*** = ***p_i−1_***.
5. Repeat steps 2 to 4 for *i* = 1,…, *N* to collect **{*p*_1_,…, *p*_*N*_}** as a sampling from the target distribution.

In the canonical version of the MH algorithm *α* is independent of the proposal distribution *Q* because it is symmetric and cancels out of the equation. In the modified version above, *Q_m_* is unknown and, in fact, is unlikely to be symmetric. In order to proceed we assume that *Q_m_* is approximately symmetric *Q_m_****(p^*^, p_i−1_)***~*Q_m_****(p_i−1_, p^*^)***, so that we can calculate *α* as above. Because of this approximation it is still necessary, following our previously published algorithm, to apply acceptance-rejection sampling to finalize the Vpop from the plausible population.

### 4.6 Assessing Similarity Between VPs

It is desirable to examine the parameter space in the Vpop to ensure heterogeneity of the VPs (while still reproducing available data). The baseline method ensured this diversity by generating VPs independently and from different initial parameter estimates. However, this is not necessarily the case for the GA and MH methods.

To assess the diversity of a Vpop we devised a test metric *d****(p_i_, p_j_)*** which scores how similar two VPs are. By randomly sampling pairs of VPs from a given Vpop, we built up a distribution for *d* and could compare the resultant cumulative density function (CDF) for each method. The test metric *d* is simply the normalized dot-product of ***p_i_*** and ***p_j_*** after they are scaled and shifted:

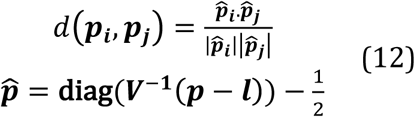

where ***V*** is a diagonal matrix such that ***v_ii_*** = ***u_i_*** — ***l_i_***. Hence, **diag(*V*^−1^(*p — l*))** uses the defined upper and lower bound for each parameter (the elements of ***u***and ***l*** respectively), to scale each parameter in ***p*** to be between 0 and 1. To ensure that *d* ∈ [—1,1] we further subtract ½ from each element. This means that, in principle, 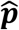 can be orientated in any direction in *m*-dimensional space (where *m* is the number of parameters). This also means that if the elements of ***p*** are sampled uniformly between the upper and lower bounds that, by symmetry, the expected value of the distribution should be zero (i.e., the CDF crosses 0.5 at *d* = 0). This is the optimal parameter set in terms of diversity, but may not be achievable given the constraints applied to the model. Conversely, if we generate Vpops from very similar parameter sets then the distribution will be right-shifted towards *d*=1.

### 4.7 Assessing Goodness of Fit

The goodness of fit (GoF) to the empirical target distribution was assessed by using the Kolmogorov-Smirnov statistic for the marginal distribution over each dimension:

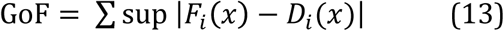

where *F_i_(x)* is the empirical cumulative distribution function for the *i^th^* model observable that we are fitting the observed cumulative distribution of the data, *D_i_(x)*. Note that for a perfect fit GoF=0, and that GoF ∈[0, *n*], where *n* is the number of distributions being fitted.

### 4.8 Source Code and Simulations

All algorithms were implemented in MATLAB 2016b (v9.1) using the Global Optimization Toolbox (v3.4.1) where a pre-packaged routine was available (e.g., GA, SA). The ODE model was implemented in SimBiology (v5.5). The K-S Test for GoF utilizes MATLAB’s Statistics and Machine Learning Toolbox (v11.0). The full source code is available for download from a GitHub repository (Rieger and Allen 2017). The source code utilizes three independent packages from MathWorks File Exchange (Johnson 2004, Jos 2016, Dorn 2017).

All simulations were performed sequentially on a MacBook Pro with a 2.9 GHz Intel Core i7 processor and 16 GB of RAM. For each choice of algorithm and pre-set number of PPs, simulations were repeated five times and the mean and standard deviation calculated.

## 5 Results

### 5.1 Comparing the various algorithms

To compare the three proposed algorithms (NSA, GA and MH) we evaluated four metrics to measure performance of the algorithm compared to the original SA method. For each algorithm, we evaluated:

1. **Efficiency:** How many PPs were needed to achieve a certain GoF of the final Vpop to the observables?
2. **Computational Cost:** How fast was the generation of PPs and VPs?
3. **Diversity:** Are the VPs parametrically similar or do they maintain the parametric heterogeneity of the PPs?
4. **Convergence:** Do the methods benefit from the acceptance/rejection step or can a Vpop be generated directly?

### 5.2 Comparison of algorithms for efficiency of yield

For each algorithm, we targeted generation of between 100 and 7,500 total PPs; those PPs were then converted into VPs through the acceptance/rejection algorithm. The GoF of the resulting Vpop was calculated as discussed in Methods. By comparing the GoF achieved for the Vpops with varying PPs (Figure 2) we find that as the number of PPs**⟶** 5,000+, all of the algorithms generated essentially indistinguishable GoFs for the final Vpop (albeit with different VPs in each Vpop). However, the three new algorithms were more efficient than the original SA method, especially when the number of PPs < 1,000. In fact, Vpops generated with as few as 100 PPs could have similar fits to the observable data as the SA method with 500+ PPs.

**Figure 2.**
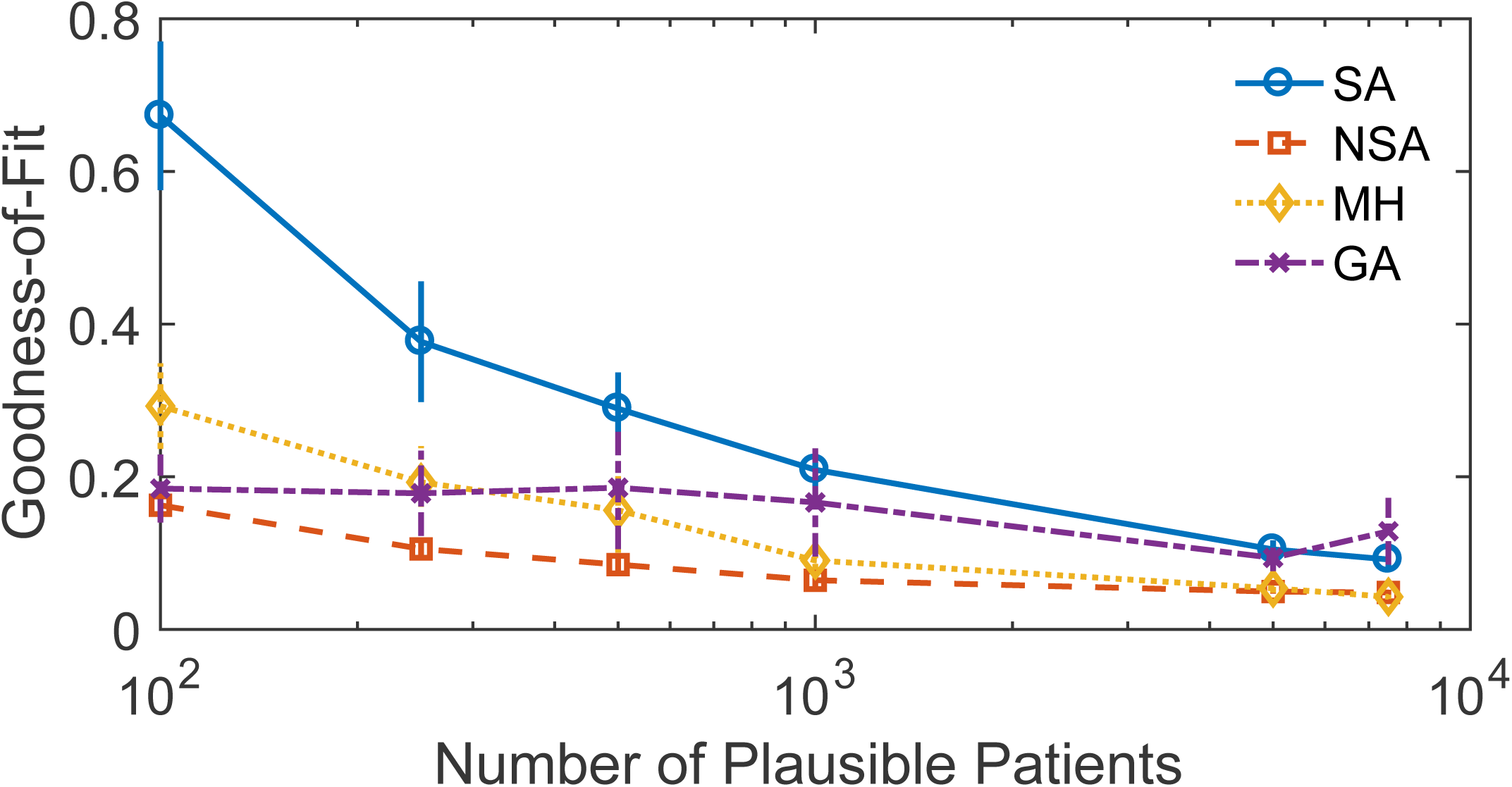
Goodness-of-fit vs. number of plausible patients. Goodness-of-fit (lower number is better) of the final virtual population vs. the number of plausible patients generated for each method. Shown is mean±standard deviation of 5 runs. SA = simulated annealing (blue circles), NSA = nested simulated annealing (red squares), MH = Metropolis-Hastings (yellow diamonds), GA = genetic algorithm (purple x’s).

### 5.3 Comparison of algorithms for computational cost

Even if an algorithm can generate the same GoF through far fewer PPs than the original SA algorithm, this does not necessarily mean the process was computationally more efficient. We further compared each method based on the clock time (evaluated via MATLAB’s *tic/toc* functions) required to generate a Vpop from 7,500 PPs (Figure 3). While the NSA method was arguably superior based on yield, this algorithm required approximately the same amount of time to execute as the SA method. Based on time, the MH and GA were the fastest algorithms and the SA remains among the least efficient.

**Figure 3.**
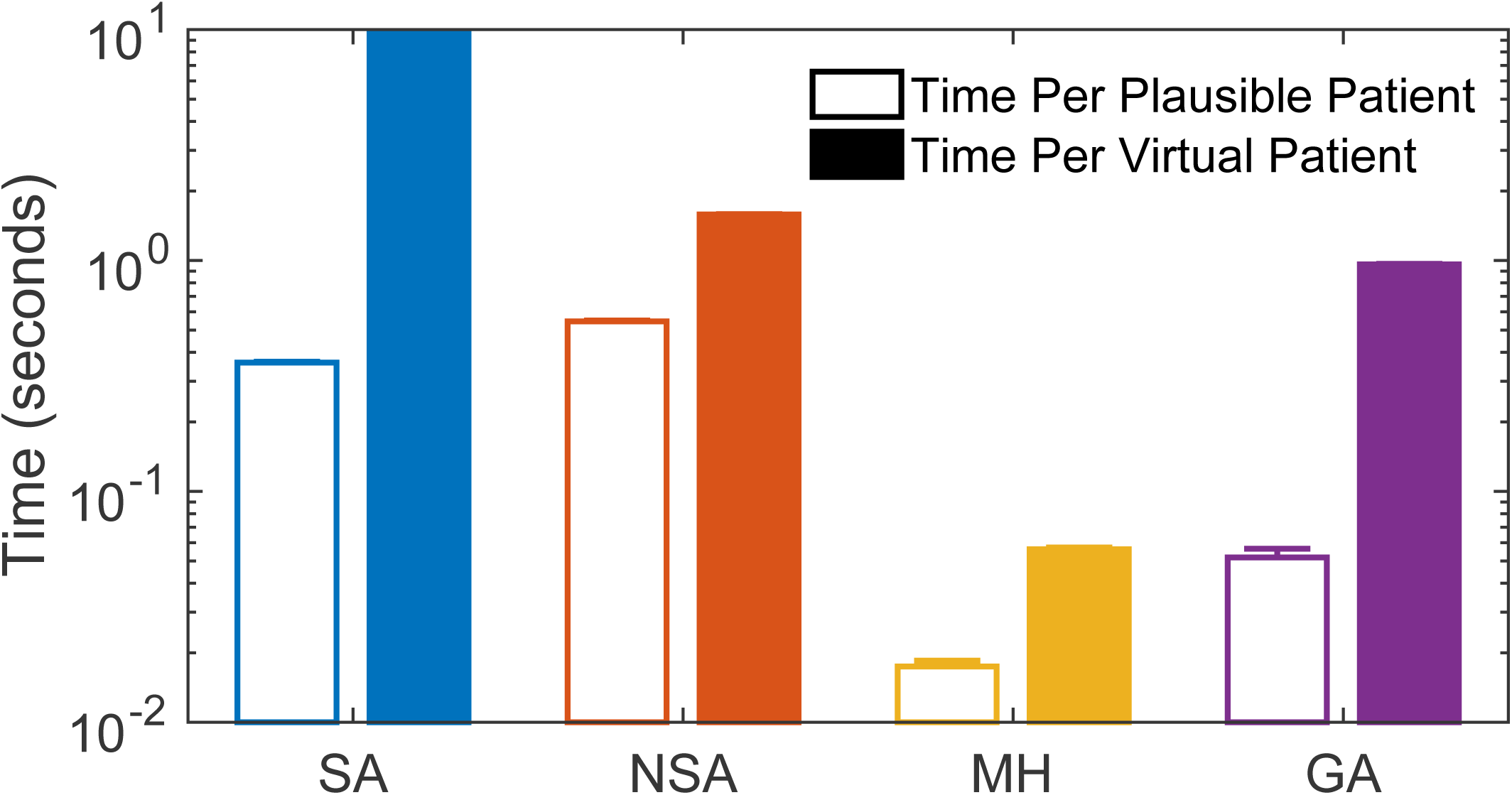
Comparison of VP and PP generation time for each method. Comparison of the time/plausible patient (open bars) or time/virtual patient (filled bars) for each method. For each method, ~7,500 plausible patients were generated and then a virtual population was selected from those plausible patients. Shown is mean±standard deviation of 5 runs. Time was calculated via the functions *tic/toc* in MATLAB. SA = simulated annealing (blue), NSA = nested simulated annealing (red), MH = Metropolis-Hastings (yellow), GA = genetic algorithm (purple).

### 5.4 Comparison of algorithms for parametric diversity of the final Vpop

An advantage of the SA algorithm is its ability to generate Vpops that maintain most of the parametric diversity of the original PPs (Supplementary Figure S2-3). This diversity in the Vpop is an essential feature for QSP models since they are often utilized in simulation of clinical trials involving novel therapies. If the underlying parameters of VPs are highly similar/correlated, clinical trial simulations performed with the Vpop may incorrectly predict a very narrow range of therapeutic response. Therefore, we need to ensure that as we introduce new algorithms, we do not trade parametric diversity for computational gains. We measured the diversity of the Vpops generated by each algorithm by uniformly sampling pairs of VPs and calculating the dot product between each set of parameters (see Methods). As a reference point/positive control, we included a set of uniformly, randomly generated model parameters (Figure 4). The closer each algorithm’s final VP parameter distribution is to the random reference, the more diverse we considered the set of VPs in the Vpop. For this criterion, the SA method was found to have the most diversity, indistinguishably followed by NSA and MH. The GA method showed distinct rightward shifts in its distribution, indicating that fewer independent parameter sets were identified in the generation of the Vpops. Supplementary figures show the violin plot for each method for both the PPs and VPs for a single iteration (Supplementary Figures S3, S5, S7, and S9).

**Figure 4.**
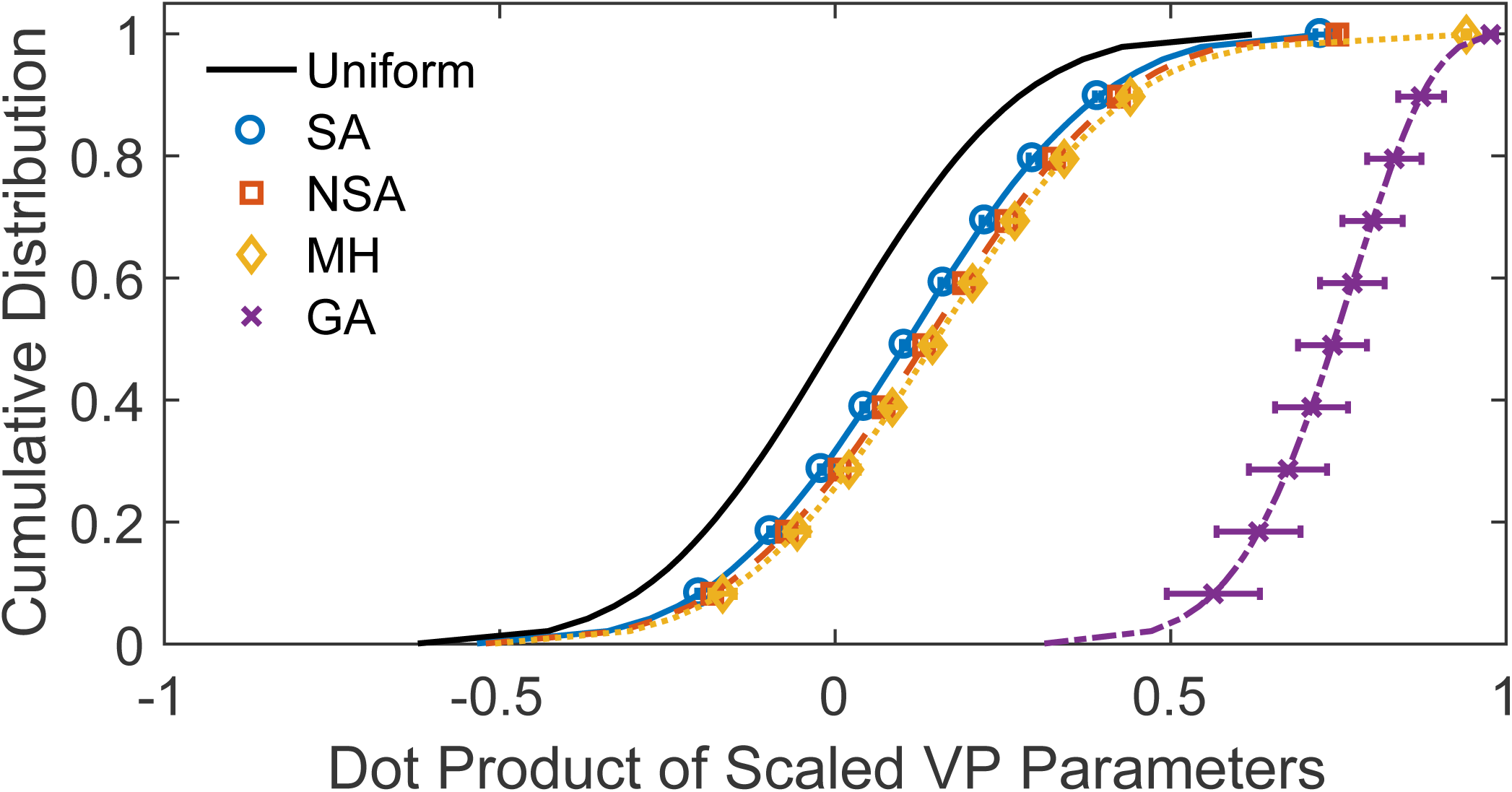
Diversity of VPs in final Vpops. Cumulative distribution vs. the dot-product of the vector of virtual patients (VPs) parameters to assess the diversity of parameter values in the virtual population. For each method, 50,000 dot-products of randomly chosen VPs were calculated and the cumulative distribution plotted. Shown is mean±standard deviation of 5 runs. As a positive control, a set of parameters from a uniform distribution was generated (solid, black line). Distributions closer to the uniform random control indicate a more diverse set of VPs. Distributions skewed towards the right indicates a more uniform set of VPs. SA = simulated annealing (blue circles, solid), NSA = nested simulated annealing (red squares, dashed), MH = Metropolis-Hastings (yellow diamonds, dotted), GA = genetic algorithm (purple x’s, dashed).

### 5.5 Convergence of the algorithms to the data

In contrast to the original SA method, each of the methods tested use information about the desired population distribution and thus requires fewer PPs to achieve an acceptable fit for the Vpop (Figure 2). To evaluate if the final selection step was still required as part of the algorithm workflow, we compared the GoF for each of the methods before and after the acceptance/rejection step, starting with ~ 7,500 PPs in each case (Figure 5). As expected, the largest improvement in Vpop fit from the PPs to VPs was for the SA method; however, all of the methods showed at least a 3-fold improvement in GoF through the final selection step. The NSA method showed the best initial fit for its PPs to the NHANES data, reasonably reproducing both the 1-dimensional and 2-dimensional histograms before the selection step (Figure S3A-F). These fits were approximately equivalent to the final Vpop fit for the SA method starting with 1,000 PPs.

**Figure 5.**
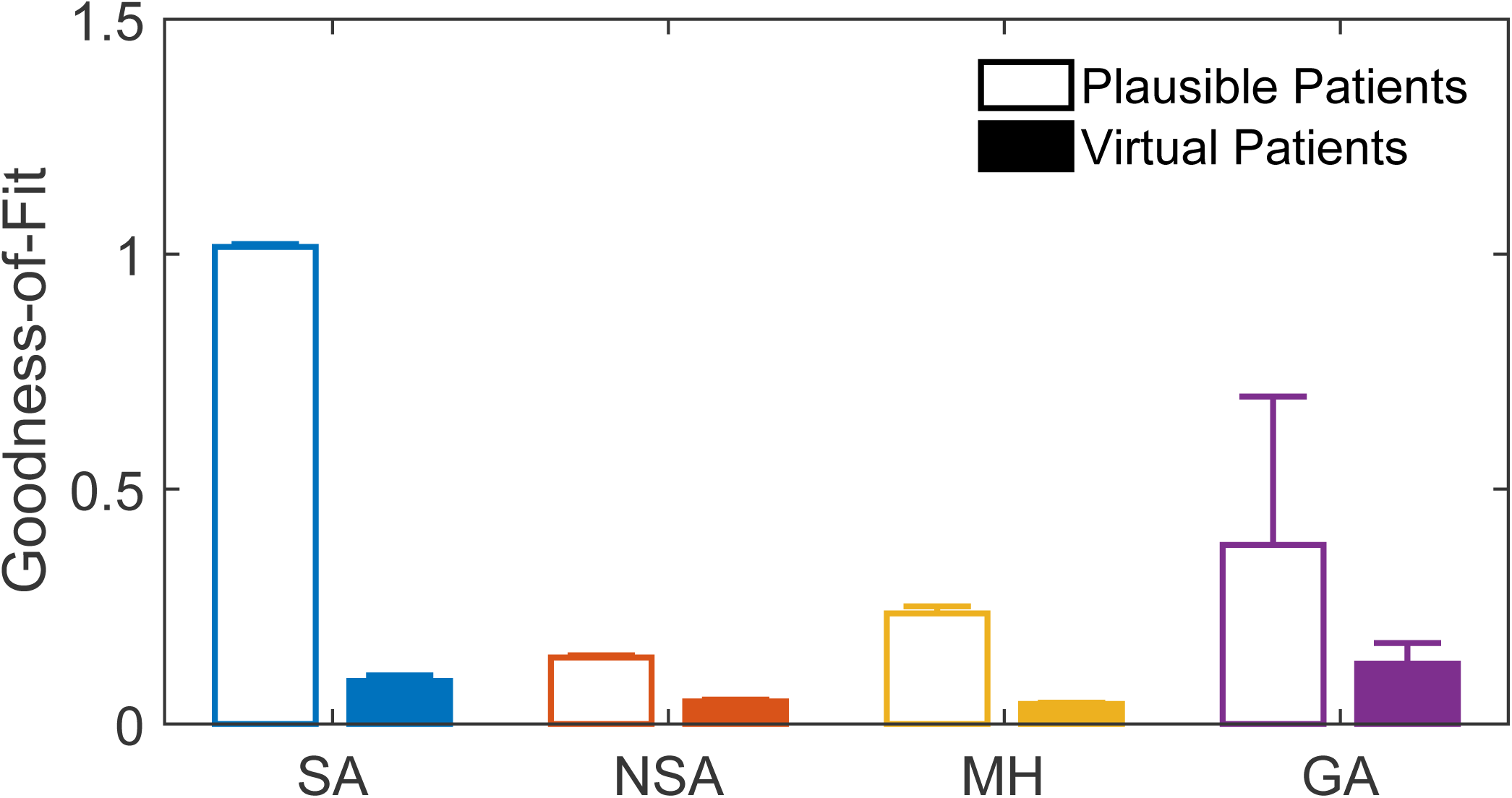
Efficiency of the acceptance/rejection algorithm. Improvement of goodness-of-fit (lower number is better) from the plausible patients (open bars) **⟶** virtual patients (filled bars) for each method starting from ~ 7,500 plausible patients. Shown is mean±standard deviation of 5 runs. SA = simulated annealing (blue), NSA = nested simulated annealing (red), MH = Metropolis-Hastings (yellow), GA = genetic algorithm (purple).

## 6 Discussion

By design, most quantitative systems pharmacology models are not identifiable from available data. While it would be desirable to have parameters well determined and characterized, the uncertainty in parameter values in QSP models often reflects our current knowledge (or lack thereof) of human (patho)-physiology. Therefore, with this perspective, we can use these models in a hypothesis-generation/testing mode to explore how these knowledge gaps translate to uncertainties in clinical outcomes and clinical trial design. In our opinion, the most thorough and robust way of doing this is by generating the most diverse Vpops given the available data.

Exploring the parameter space of under-determined quantitative systems pharmacology models remains a challenge but it is essential for robust predictions of safety and efficacy for novel compounds. Here we presented three methods for generating diverse parameter sets in a QSP model. While this exercise is by no means an exhaustive exploration of global optimization techniques, each algorithm improved at least one of the testing metrics compared to our previously published SA method. The seemingly simple question of which method is “the best” cannot be definitively answered here but it is important to be aware of the pros and cons for each and potential steps to improve performance.

We previously discussed the advantages and disadvantages of using the SA method for this application (Allen, Rieger et al. 2016). In comparison to other methods tested it was the slowest (or tied with NSA) and required the most PPs for a quality fit. Conversely, it also generated the most diverse Vpop with the fewest imposed correlations. The ease of implementation is also an advantage for SA. As implemented, the algorithm required no prior knowledge of the final Vpop distribution and there was a minimal set of tuning parameters required, most of which were default MATLAB options (i.e., no arbitrary decision about number of ellipsoids, number of generations). As such, the algorithm remains relevant as a “first try” for generating VPs. Furthermore, it is the only method that can be run (at least in part) without prior knowledge of the target distribution. This relaxed requirement can make it an attractive choice for pre-computing plausible populations or for exploring how the parameter space relates to the model output (for example, identifying parameters in the model that can give rise to sub-populations of interest).

The NSA approach iterates on the SA method by utilizing prior knowledge of the final parameter distribution and forcing the algorithm to regions with the most desired patient density. This method is likely most efficient when the target distribution is approximately multivariate normal. Fortunately, for the example here, the target distribution was well approximated as a multivariate lognormal. The viability of this method for more eccentric or bimodal distributions would need to be studied by applying it to other case studies. However, such distributions are less commonly observed in clinical trials. For our case study, the computational cost to generate a plausible population was comparable between the NSA and SA methods; however, the NSA approach demonstrated vastly improved yield (VP per PP), which facilitated an overall more efficient Vpop generation process. The algorithm was as good, or better, than the other methods tested for direct generation (pre-selection/rejection) of a reasonable Vpop, without the imposed correlations found in Vpops generated by GA. While essentially the same as the SA method to implement, there is a problem-specific choice for the number of ellipsoids to use. Here we used five regions, regardless of the number of PPs being generated. Fine-tuning based on the model/number of PPs may potentially improve performance.

MH is unique amongst the approaches we tried in that it is a Markov Chain, which should imply some degree of correlation between the PPs. The advantage of this technique was an increase in speed and compared to the two methods based on SA; however, there also was a right-shift in the dot product cumulative distribution, implying a slightly less diverse final population. Methods have been published to attempt to reduce this correlation (Santoso, Phoon et al. 2011) and to improve performance in higher dimensions (Betancourt 2017), but we chose to evaluate only the common form of the algorithm and to leave further exploration for future improvements. While straightforward to implement, MH requires the choice of a proposal distribution. As noted in the Methods, because we indirectly sample the distribution of the observables by first sampling the parameter space and then generate the observables through model simulation we do not have direct control over the choice of a proposal distribution. The implications of this for direct convergence of this method will depend on the symmetry of the observable distribution (induced by the parameter sampling) around every point on the Markov chain. In this case, the approximation we assumed (see Methods) appeared to hold sufficiently for the MH algorithm to approximate the empirical distributions. However, the final Vpop was improved by acceptance/rejection sampling.

The GA was very similar to MH in that, compared to the SA-based methods, a 10-fold improvement in computational speed was achieved at the cost of some lost heterogeneity in the final Vpop. The GA is easy to implement for QSP models and the supplied routine with MATLAB’s Global Optimization Toolbox was sufficient for our purposes. GA requires some problem-specific decisions, which may affect overall performance; for example, the size of the population, number of generations, and mutation rate can be adjusted as needed.

The curse of under-determined models has led to a long history of using different global optimization techniques for generating parameter sets within the bounds of the data (van de Pas, Woutersen et al. 2012, Gadkar, Budha et al. 2014, Hallow, Lo et al. 2014). Use of global optimization techniques often feels like more of an art than a science due to how problem-specific their application can be. For this reason, we examined several algorithms with different approaches for exploring constrained, multidimensional parameter spaces. Requiring only minimal tuning, each of these algorithms successfully explored the range of our 23-dimensional parameter space and generated reasonable PPs. The choice of algorithm to use for a new problem, particularly one with higher dimensions and a less Gaussian set of observations, will need to be evaluated on a case-by-case basis. For example, for models that are slow to simulate the most constraining factor is computational cost. In this case the MH or GA approaches may be the most successful; however, as we have shown, without adaptation these methods may come at a cost of diversity in the final Vpop.

We hope that the results presented here will provide a guide to selection and implementation of these algorithms to facilitate the generation of robust Vpops in mathematical models of (patho)-physiology.

## 7 Author contributions

RJA conceived of the original algorithm. CJM, RJA, and TRR conceived of the updated algorithm objectives. AG, GWC, LB, RW, YC1, YC2, and YL selected and coded the new algorithms and performed the initial proof-of-concept testing. CJM, RE, HTB, RJA, and TRR supervised and advised the initial work. RJA, RE, RW, TRR, HTB, and CJM drafted the manuscript. All authors reviewed, revised, and approved of the final manuscript.

## 10 Acknowledgements

The authors thank Ilse Ipsen and Thomas Gehrmann for organizing the 2016 SAMSI Workshop at North Carolina State University, which initially facilitated this collaboration.

## Abbreviations

GA: Genetic Algorithm
GoF: Goodness of Fit
HDL_c_: High Density Lipoprotein Cholesterol
LDL_c_: Low Density Lipoprotein Cholesterol
MH: Metropolis-Hastings
NHANES: National Health And Nutrition Examination Survey
NSA: Nested Simulated Annealing
SA: Simulated Annealing
PP: Plausible Patient
QSP: Quantitative Systems Pharmacology
TC: Total Cholesterol
VP: Virtual Patient
Vpop: Virtual Population

